# Identification and predictive machine learning models construction of gut microbiota associated with lymph node metastasis in colorectal cancer

**DOI:** 10.1101/2024.10.10.617644

**Authors:** Yongzhi Wu, Xiaoliang Huang, Yongqi Huang, Zigui Huang, Chuanbin Chen, Mingjian Qin, Zhen Wang, Fuhai He, Binzhe Tang, Chenyan Long, Xianwei Mo, Jungang Liu, Weizhong Tang

## Abstract

This study focuses on the significant role between gut microbiota and lymph node metastasis (LNM) in colorectal cancer (CRC). By conducting 16S rRNA sequencing on fecal samples from 147 CRC patients and combining it with the linear discriminant analysis effect size (LEfSe) algorithm, we successfully identified significant differences in the gut microbiota between patients with LNM and those with none lymph node metastasis (NLNM). Furthermore, using transcriptome data from 9 CRC patients, we constructed an immune cell infiltration matrix to deeply explore the biological functions associated with LNM. Eventually, using the characteristics of the gut microbiota associated with LNM, we developed random forest (RF) and multilayer perceptron (MLP) machine learning models to predict the lymph node metastasis status of CRC patients. We identified 21 differentially abundant gut microbes between the two groups, among which *Bacteroides plebeius*, significantly enriched in LNM group, is closely related to the upregulation of resting NK cells and chemokine CXCL8 expression, and this bacterial species is also positively correlated with the enhancement of fructose metabolism. The RF and MLP models constructed based on the LNM-associated gut microbiota showed good predictive efficacy in predicting LNM status in CRC. This study reveals that *Bacteroides plebeius* may play an important role in the progression of CRC, with its mechanism potentially involving changes in immune modulation and metabolic pathways. The classification model constructed based on gut microbiota characteristics can predict LNM status of CRC, providing a new perspective for personalized and precision treatment of CRC patients.

**Importance:** This study highlights the pivotal role of gut microbiota in lymph node metastasis (LNM) of CRC, identifying key microbial differences between LNM and NLNM patients. Our findings implicate *Bacteroides plebeius* in CRC progression via immune modulation and metabolic alterations. Moreover, machine learning models based on gut microbiota predict LNM status accurately, offering a novel approach for personalized CRC treatment.

## 1. Introduction

Colorectal cancer (CRC) ranks as the third most common malignant tumor and the second leading cause of cancer-related deaths worldwide, with continued attention on its prevention and treatment^[1]^. However, there are still over 1.9 million new cases and 900,000 deaths annually. In China, due to rapid economic development in recent years, lifestyle changes^[2]^ such as physical inactivity, smoking, alcohol consumption, and dietary habits characterized by high fat, high protein, and low fiber have led to a steady increase in the incidence of CRC. It is the second most common type of cancer among the top five in China^[3]^. Furthermore, due to the adverse side effects associated with treatment and the poor prognosis of local recurrence and distant metastasis^[4]^, the mortality rate of CRC remains high, imposing a heavy burden on patients and society.

After years of research, it has been found that CRC is a multifactorial disease caused by a variety of genetic, external, and internal environmental factors leading to tumor formation and disease progression^[5]^. A growing body of evidence suggests that the abundance and structure of gut microbiota and their metabolites, as an internal environmental factor, play an immeasurable role. The human gut harbors more than 1,000 different species and over 7,000 different strains of bacteria^[6]^, which are not only involved in the digestive process of the human body but also maintain immune system balance and regulate metabolic functions. As the largest microbial ecosystem in the human body, the complexity and diversity of the gut microbiota are beyond imagination. These microorganisms are closely linked to host health through various means such as producing a variety of metabolic products, participating in nutrient absorption and metabolism, and affecting the function of the intestinal barrier and immune response^[7]^. Studies have shown that *Fusobacterium nucleatum* can promote the occurrence of CRC by activating and inhibiting apoptosis through microRNA-mediated Toll-like receptor 2 (TLR2)/Toll-like receptor 4 (TLR4) signaling pathways^[8]^. Research by Wu et al. confirmed^[9]^ that in mouse models, oral administration of *Fusobacterium* can change the composition of the gut microbiota and promote cancer development by activating the AMPK signaling pathway, thereby inhibiting the production of butyrate. Certain probiotics in the gut, such as *Bifidobacterium* and *Lactobacillus*, can maintain the balance of gut microbiota^[10]^ by producing short-chain fatty acids (SCFA), thereby regulating host immune responses, inducing cancer cell apoptosis, and inhibiting tumor cell proliferation^[11]^. It can be seen that in CRC, a malignant tumor that poses a serious threat to human health, the role of gut microbiota is increasingly valued.

Lymph node metastasis refers to the phenomenon where infiltrating tumor cells penetrate the walls of lymphatic vessels, detach and are carried to the regional lymph nodes by lymphatic fluid, and grow into the same type of tumor centered around these nodes^[12, 13]^. Lymph node involvement has significant guiding implications for the treatment and prognosis of colorectal cancer; a positive status can identify patients who need adjuvant chemotherapy^[14]^, and a negative status is also an important independent factor for the prognosis of patients with high microsatellite instability (MSI-H)^[15]^. The College of American Pathologists (CAP) and the American Joint Committee on Cancer (AJCC) believe that at least 12 lymph nodes^[16]^ need to be examined during surgical resection to ensure accurate staging. In CRC, the presence of cancer cells in the tumor-draining lymph nodes is defined as stage III disease, but some models consider lymph node metastasis as a harbinger of distant metastasis^[17]^, and lymph node and distant metastases do indeed often share a common origin^[18]^. Therefore, in recent years, more and more research has been dedicated to predicting lymph node status. Some studies have used patient age, gender, tumor size, location, morphology, vascular invasion, histological grading, and other data to establish a machine learning artificial neural network to identify T1 colorectal tumors at risk of lymph node metastasis^[19]^. Xia et al. constructed a diagnostic model for lymph nodes in rectal cancer patients by utilizing preoperative MRI data combined with postoperative patient pathological information^[20]^. However, to date, there is no complete and accurate predictive model to precisely identify the lymph node metastasis status.

This study collected preoperative fecal samples from 147 CRC patients and postoperative tumor samples from 9 patients, performing 16S rRNA sequencing and transcriptome sequencing respectively, to explore the differential gut microbiota and biological functions between LNM-CRC and NLNM-CRC patients, identify biomarkers and corresponding biological mechanisms associated with LNM risk, and construct a machine learning diagnostic model combined with characteristic microbiota to accurately predict the LNM status of CRC patients, providing a new theoretical perspective for clinical treatment decisions.

## 2. Result

### 2.1 Clinical data and statistical characteristics of CRC patients enrolled in the study

Researchers collected 198 fecal samples from CRC patients who met the study criteria for 16S rRNA sequencing analysis. A total of 147 patients with complete LNM status data were selected. They were then divided into two groups based on lymph node negativity and positivity: the non-lymph node metastasis group (NLNM) with 69 individuals, and the lymph node metastasis group (LNM) with 78 individuals. The detailed clinical characteristics of the patients are shown in Table.1. There were no significant differences in average age, gender distribution, BMI, tumor localization, tumor volumes, and MMR status between the two groups (P > 0.05), indicating that the baseline characteristics of the CRC patients in this study were balanced and comparable.

### 2.2 Comparison of microbiome diversity between LNM and NLNM

In order to assess potential differences in gut microbiome diversity between LNM and NLNM, we conducted an analysis of α-diversity. Fig.1A shows that the six α-diversity indices studied did not exhibit significant statistical differences between groups (P > 0.5). The gut microbiota of the two groups were screened and visualized as shown in the Venn diagram in Fig.1B, revealing certain characteristics and commonalities among samples from different groups. The PLS-DA analysis depicted in Fig.1C further successfully distinguished the gut microbiota of the LNM group from that of the NLNM group into separate clusters. These results indicated that while there were no significant differences in the overall diversity levels of the gut microbiota between groups, distinct compositional differences persisted in the gut microbiota between the LNM and NLNM groups.

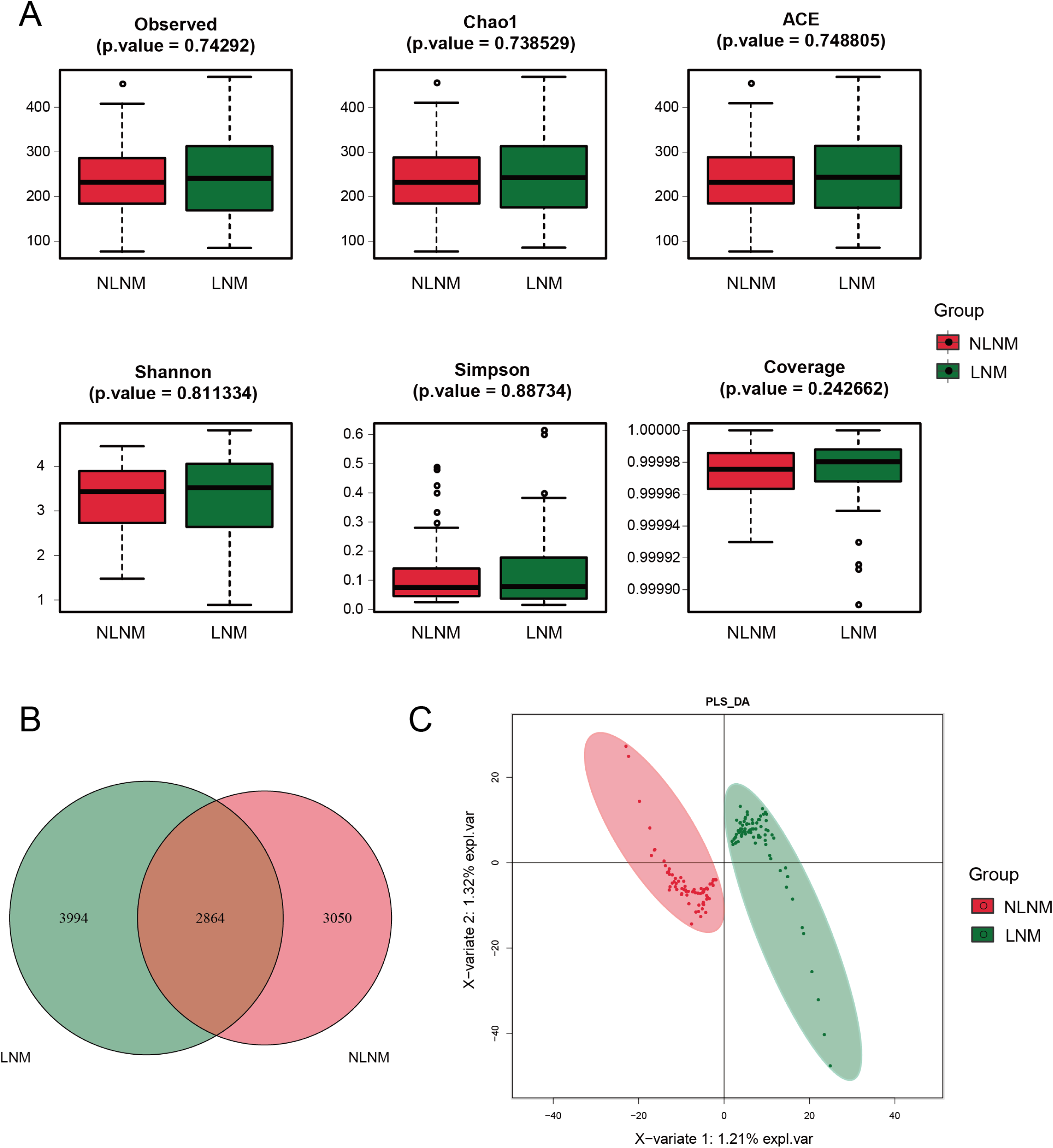

### 2.3 Exploration of LNM-associated gut microbiota

In order to identify the differentially abundant gut bacteria between the LNM and NLNM and to find potential LNM-associated gut microbiota biomarkers, we conducted a linear discriminant analysis effect size (LEfSe) on the gut microbiota of the two groups (Fig.2A-B and Table.S1). The LDA bar chart in Figure 2B illustrates the LDA scores from the LEfSe analysis of differentially abundant gut microbiota between the two groups (scores are log10-transformed), with higher LDA scores indicating stronger discrimination between the groups. It was found that the abundance of 21 species showed statistically significant differences, with 12 species significantly higher in LNM group than in NLNM group. Furthermore, to further investigate the interactions between the differential gut microbiota that were significantly elevated in the LNM and NLNM groups, we plotted the correlation graph of the dominant gut microbiota in the two groups (Fig.2C). Among them, the dominant bacteria in the NLNM group, *o Actinomycetales.f Corynebacteriaceae* and *f Corynebacteriaceae.g Corynebacterium*, and the dominant bacteria in the LNM group, *g Vagococcus.s Vagococcus_teuberi*, were the three bacteria most closely connected to other nodes. This indicates that these three bacteria have the closest correlation with other dominant bacteria. Additionally, we found that *g Howardella.s uncultured_organism* in the LNM group showed a significant negative correlation with *o Actinomycetales.f Corynebacteriaceae* and *f Corynebacteriaceae.g Corynebacterium* in the NLNM group. Based on these results, we speculated that there may be potential competitive relationships between the dominant microbial communities in the two groups.

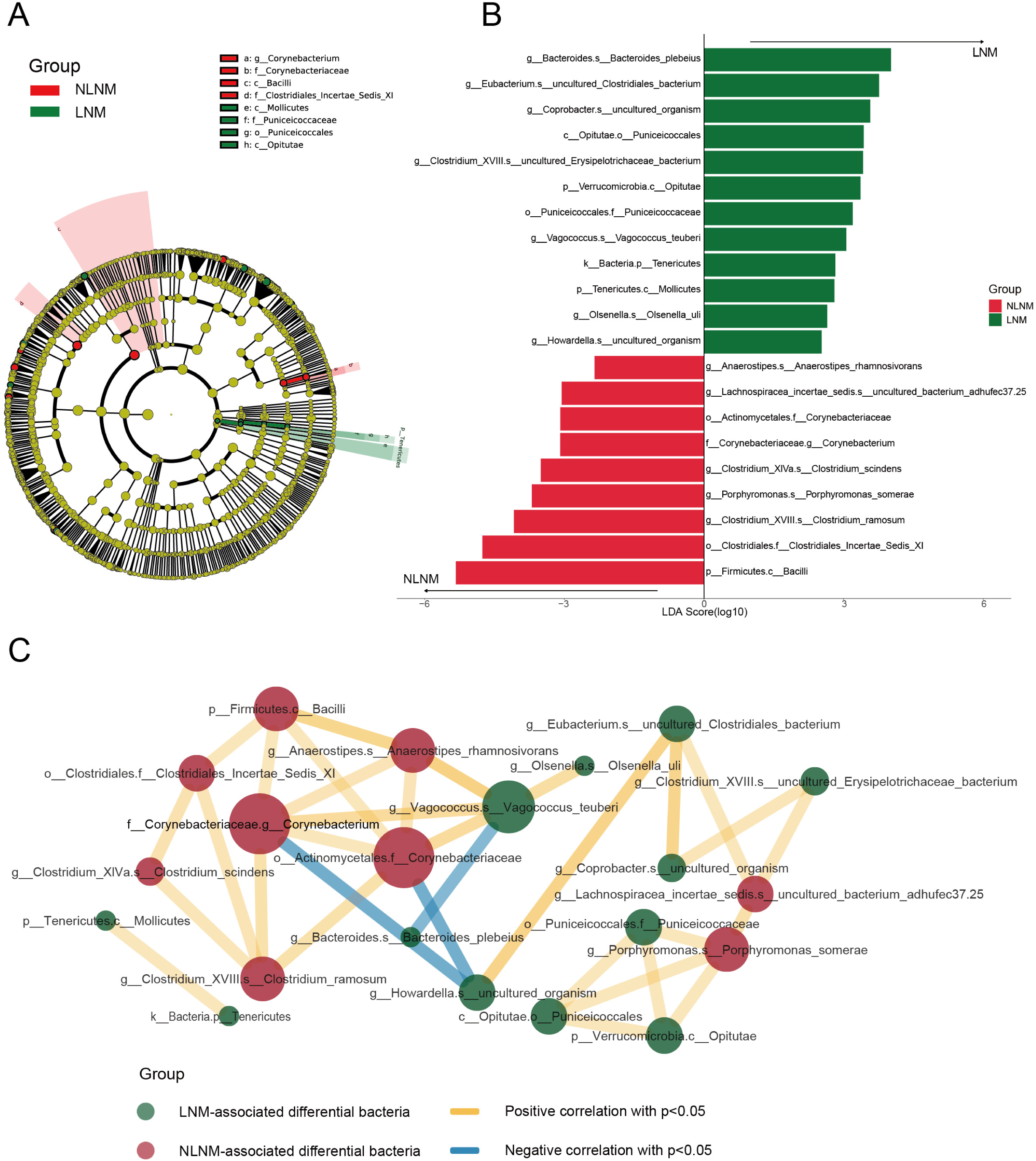

### 2.4 Biological functional prediction of gut microbiota in LNM and NLNM

To explore the biological pathways enriched by genes in the gut microbiome of colorectal cancer patients under different lymph node states, Phylogenetic Investigations of Communities by Reconstruction of Unobserved States II (PICRUSt 2) was used to predict the KEGG pathways of LNM and NLNM patients. A total of 174 different KEGG pathways were identified, among which 4 pathways showed statistically significant differences (Fig.3 and Table.S2) (P < 0.05). Specifically, there was 1 pathway with significantly higher abundance in LNM group compared to NLNM group, which is the vibrio cholerae infection pathway (P = 0.034). In NLNM group, there were 3 pathways with significantly higher abundance than in LNM group, namely the renin-angiotensin system (P = 0.016), steroid biosynthesis (P = 0.028), and endocytosis (P = 0.033). It can be inferred that CRC patients with negative and positive lymph nodes may exhibit differences in gut metabolic characteristics.

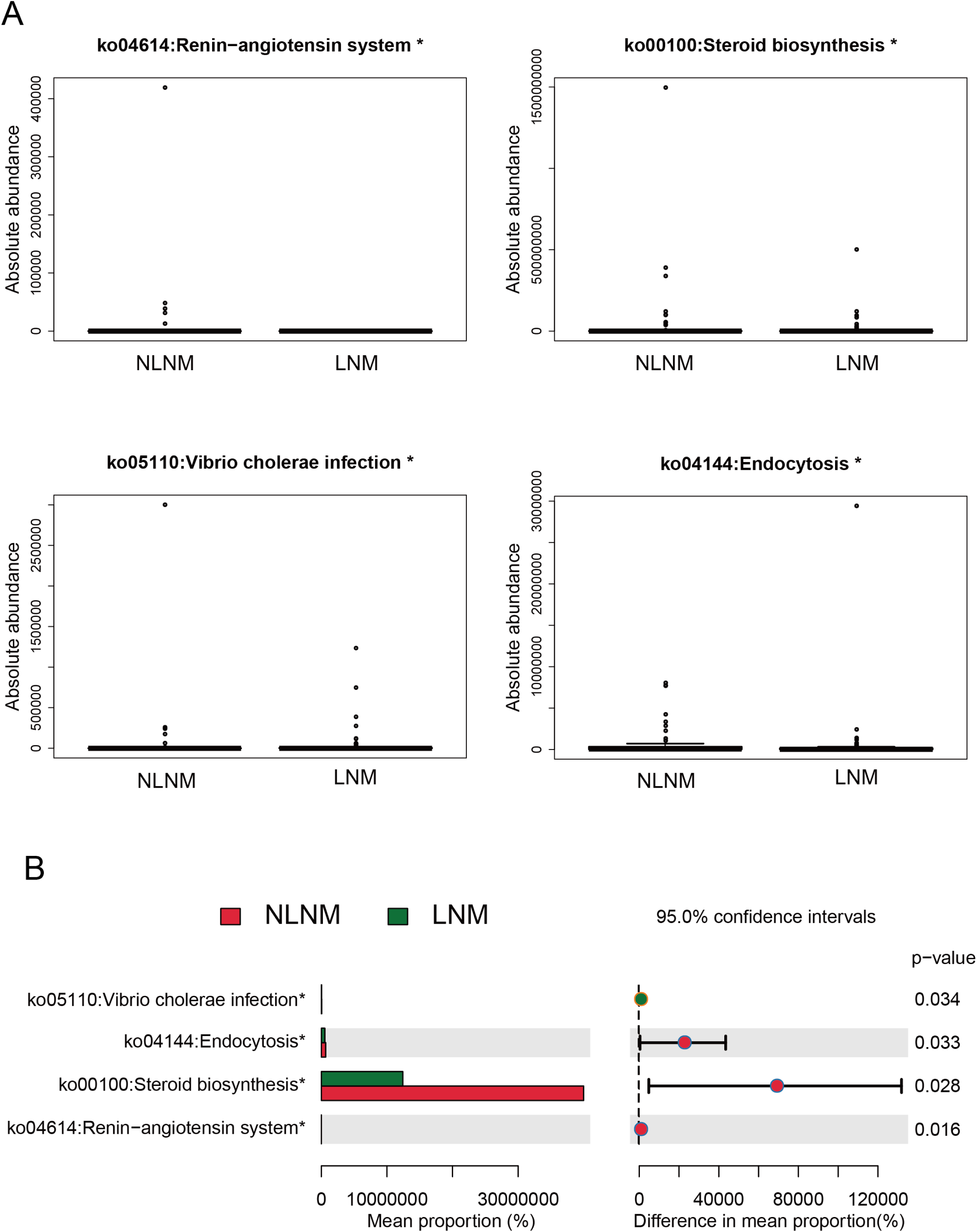

### 2.5 Correlation between LNM-related gut microbiota and tumor-infiltrating immune cells

Tumor-infiltrating immune cells play a crucial role in the tumor immune microenvironment, influencing tumor-associated immune responses, thereby inhibiting tumor growth or potentially promoting tumor metastasis and immune evasion^[21]^. Tumor-infiltrating immune cells are potential targets for cancer immunotherapy. Therefore, we utilized RNA sequencing data to investigate the composition of 22 infiltrating immune cells in 9 CRC patients (5 in NLNM group and 4 in LNM group), and displayed the abundance of immune cells in the form of a ribbon plot (Fig.4A). As shown, it illustrates the different characteristics of the immune tumor microenvironment in patients with different lymph node statuses. Overall, we can observe that the abundance of B cells memory and Macrophages M1 is lower in LNM group compared to NLNM group. Subsequently, by comparing the dominant gut microbiota of two groups with the 22 immune cells, we analyzed the correlation between LNM-associated differential microbiota and immune cells (Fig.4B-C). In NLNM group, *g Porphyromonas.s Porphyromonas_somerae* is significantly positively correlated with T cells CD4 memory activated and Monocytes. In LNM group (Fig.4C-D), *g Coprobacter.s uncultured_organism* is significantly positively correlated with Dendritic cells activated and T cells CD4 memory activated, while significantly negatively correlated with Macrophages M2. At the same time, *g Bacteroides.s Bacteroides_plebeius* is positively correlated with NK cells resting. These findings suggest that there are differences in immune cell infiltration between the LNM and NLNM groups, and that LNM-associated dominant gut microbiota in CRC patients are distinctly associated with various tumor-infiltrating immune cells, implying a potential regulatory role of LNM-associated differential gut microbiota in shaping the CRC immune microenvironment.

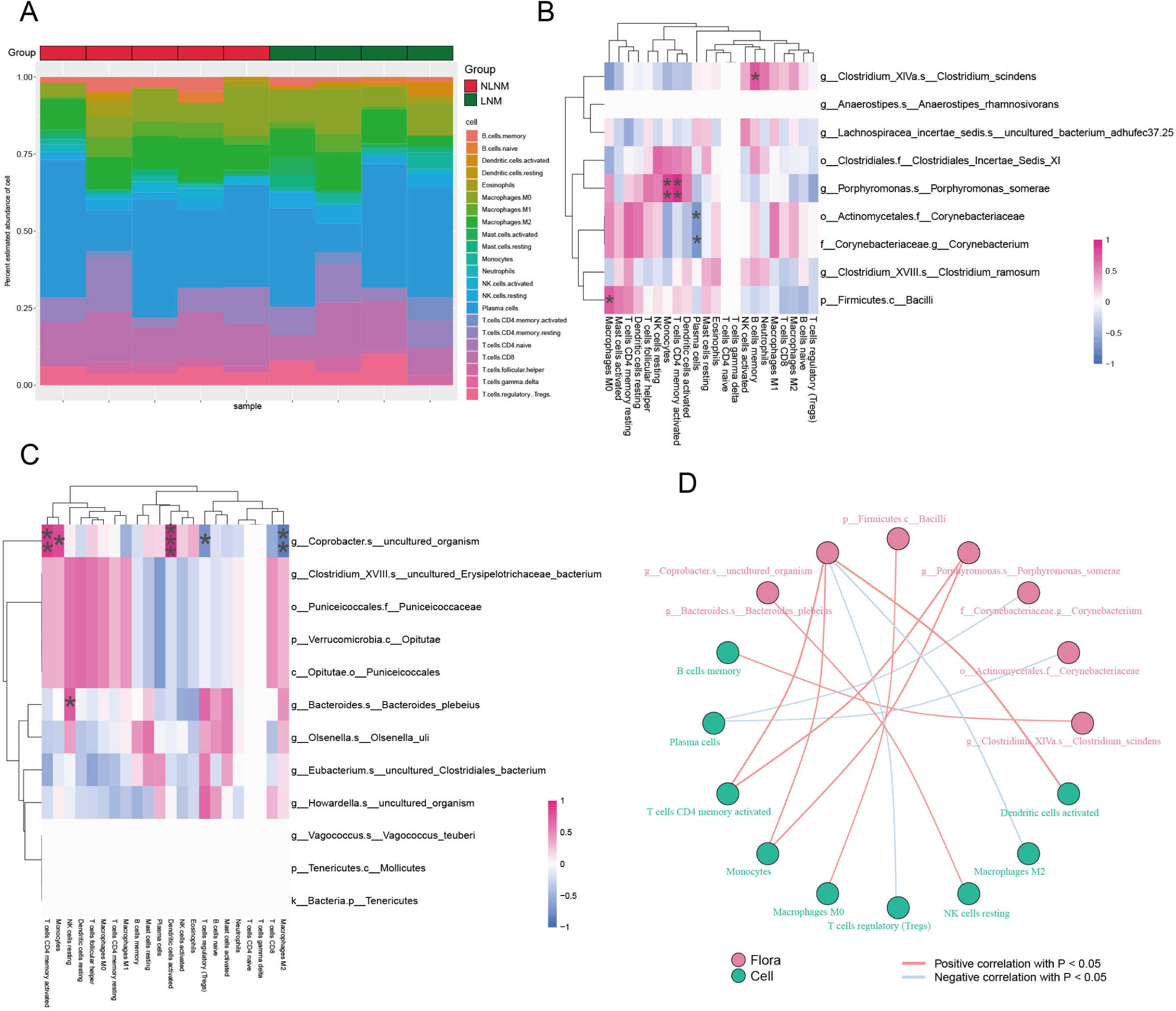

### 2.6 Relationship between LNM-related variations in gut microbiota and immune-associated genes

By analyzing the correlation between LNM-associated differential gut microbiota and common immune-related genes, we discovered a potential link between LNM-associated gut microbiota and host immunity. As shown in Fig.5, among the differential bacteria in NLNM, *g Lachnospiracea_incertae_sedis.s uncultured_bacterium_adhufec37.25*is significantly positively correlated with various immune checkpoints (VTCN1, LAG3, etc.) (Fig.5A), chemokines (CXCL12, CCL25, CCL14, CCL11, etc.) (Fig.5B), immune suppressor genes (KDR, IL10, etc.), immune activation genes (BTNL2, ENTPD1, TNFRSF17, etc.), and chemokine receptors (CCR2) (see Fig.S1). Among the differential bacteria in LNM, g Clostridium_XVIII.s uncultured_Erysipelotrichaceae_bacterium is significantly positively correlated with various immune checkpoints (KIR3DL1, LAIR1, etc.) (Fig.5C) and chemokines (CCL1, CCL7, etc.) (Fig.5D). The human immune system plays a crucial role in tumor occurrence and development. These findings suggest that the differential gut microbiota associated with LNM may influence the expression of immune-related genes.

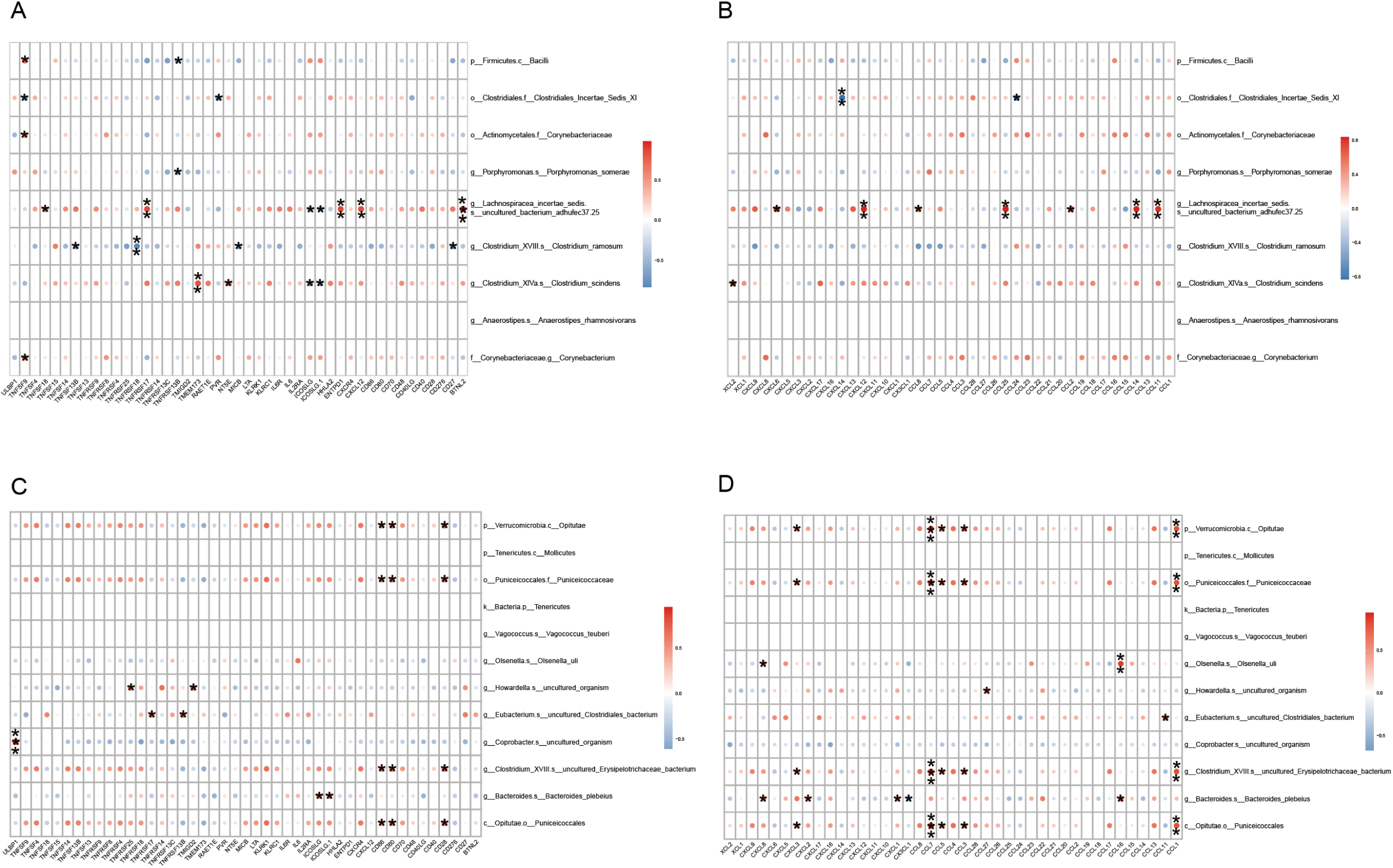

### 2.7 Identification of LNM-associated biological functional pathways and correlation of differential pathways with differential gut microbiota

In order to identify the differential biological functional pathways between LNM and NLNM and to investigate the correlation between different gut microbiota in two groups and these pathways, we utilized ssGSEA to convert the gene expression matrix obtained from RNA sequencing of tumor tissue samples from nine CRC patients and the species abundance matrix obtained from 16S rRNA sequencing of gut microbiota into corresponding score matrices. Subsequently, these matrices were analyzed through KEGG and GO analyses, with GO analysis including cellular component (CC), molecular function (MF), and biological process (BP). After performing differential analysis on the GO (Fig.6A) and KEGG (Fig.6B) pathway score matrices for the LNM and NLNM groups, in LNM group, we identified 80 significantly upregulated GO pathways [such as GOBP: regulation of sequestering of triglyceride (logFC=0.044, P=0.006) and GOBP: regulation of low-density lipoprotein particle clearance (logFC=0.036, P=0.002)]. In NLNM group, a total of 32 significantly upregulated GO pathways were identified [such as: GOBP: cytidine to uridine editing (logFC=0.047, P=0.004) and GOMF: vinculin binding (logFC=0.031, P=0.009)]. However, no significant KEGG differential pathways were found in either group (Fig.6B). The complete information of the GO enrichment items can be found in Table.S3. These findings highlight the different biological functions between LNM and NLNM. We further investigated the correlation coefficients between each group of bacteria and BP items and MF items (Table.S4), and found that there is a significant correlation between certain gut microbiota and biological pathways (Fig.6C). For example, in LNM group, *g Bacteroides.s Bacteroides_plebeius* has a statistically significant positive correlation with GOBP: fructose metabolic process (r=0.84, P=0.004).

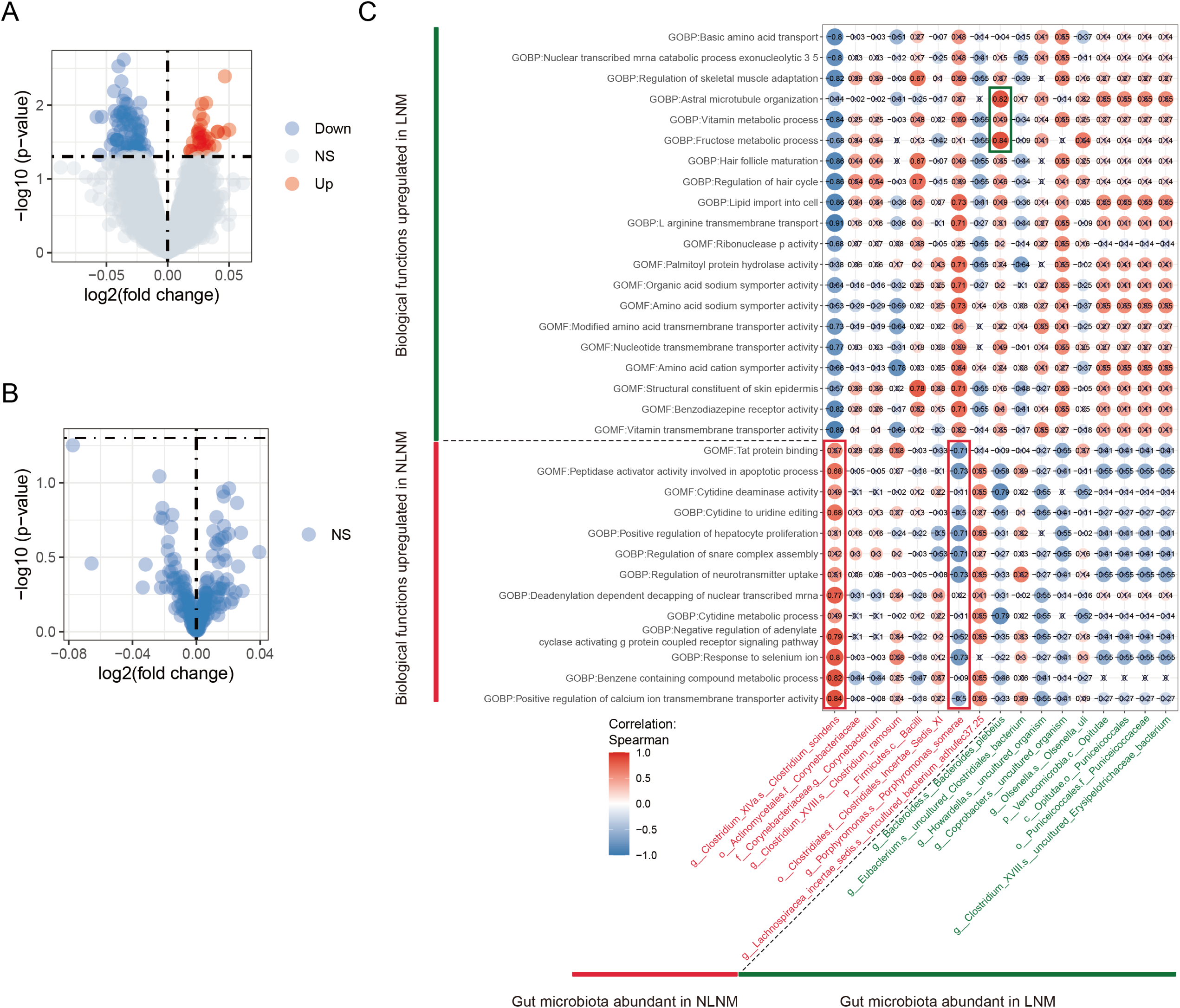

In NLNM group, *g Clostridium_XlVa.s Clostridium_scindens* has a statistically significant positive correlation with GOBP: positive regulation of calcium ion transmembrane transporter activity (r=0.84, P=0.004). These results suggest that the differential gut microbiota associated with LNM may influence the metastasis of cancer cells to lymph nodes by participating in certain potential biological functional pathways.

### 2.8 Construction of a model to predict LNM status using differential gut microbiota characterization

As shown in Fig.2B, we have identified 21 differential gut microbiota associated with LNM based on LEfSe analysis. After a detailed ranking of the importance of gut microbiota features, we retained 13 of the higher-ranking microbial features (accounting for 62%) to develop random forest (RF) and multilayer perceptron (MLP) predictive models based on LNM-related gut microbiota. The performance evaluation of the RF model showed that the area under the curve (AUC) value for the training set was 0.952, and the AUC value for the validation set was 0.796, close to 0.8 (Fig.7A). Similarly, the receiver operating characteristic (ROC) curve based on the MLP model also performed well, with an AUC value of 0.963 for the training set and an AUC value of 0.793 for the validation set (Fig.7B). This indicates that both algorithmic models have good performance in predicting whether patients have lymph node metastasis, providing a new strategy for the precise prediction of lymph node metastasis in clinical practice and the formulation of subsequent treatment plans.

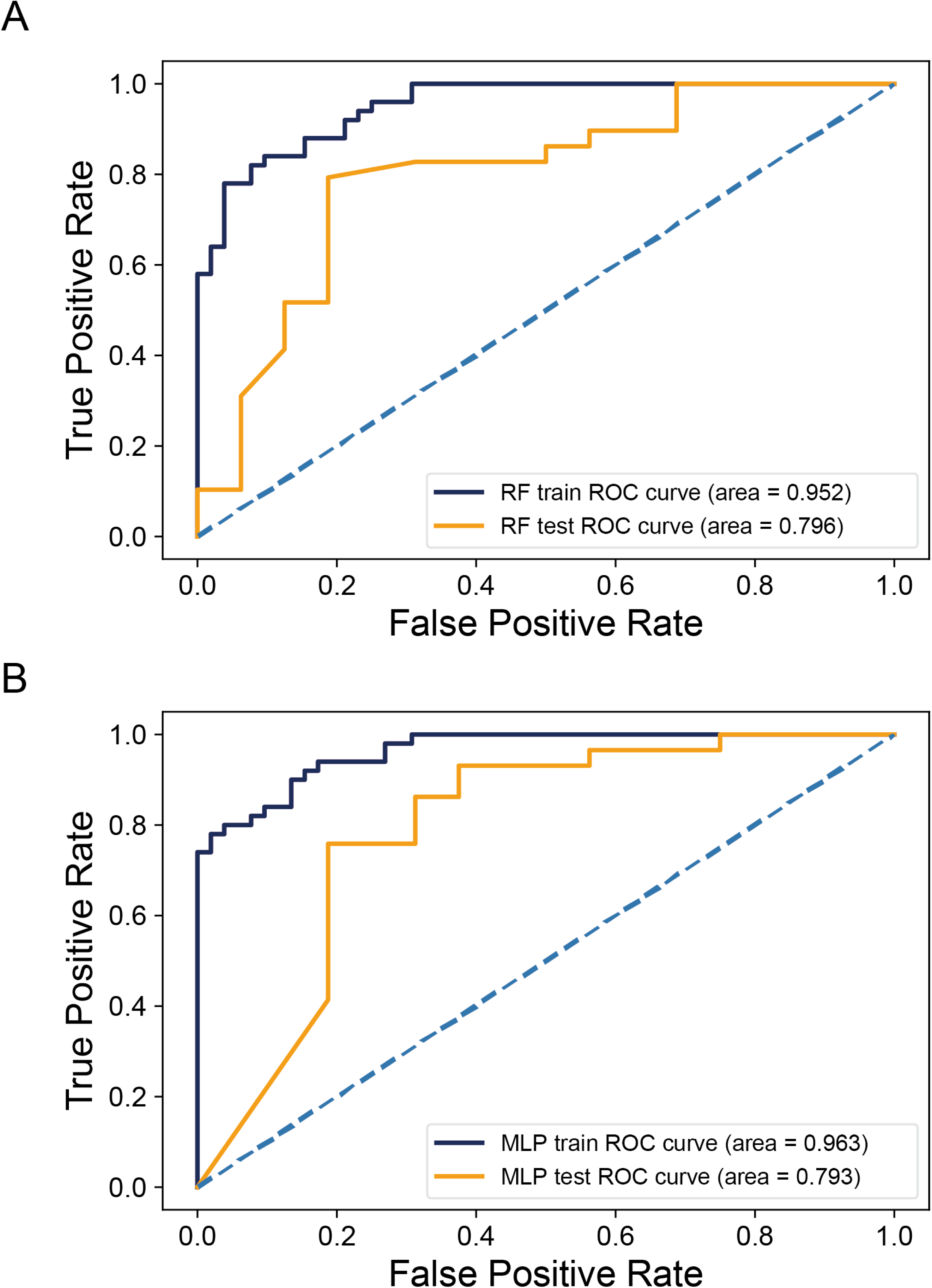

## 3. Discussion

Having employed 16S rRNA sequencing and transcriptome sequencing technologies, we meticulously analyzed the community diversity of gut microbiota and differences in dominant bacterial groups between LNM and NLNM CRC patients. We clarified the relationship between dominant bacterial groups and host immunity and deeply explored the potential biological pathways associated with LNM. After identifying differential gut microbiota, we then used these microbial features to construct a classification model to predict LNM status.

Through α-diversity analysis of fecal samples from two groups of tumor patients, we observed that these indices did not show significant differences statistically (see Fig.1A). However, other studies have found that the α-diversity of the gut microbiota in CRC patients is significantly higher than that in healthy control groups^[22]^, indicating that the occurrence and development of CRC are associated with changes in the diversity of the gut microbiota. Subsequent Venn diagram and PLS-DA analyses found that the gut microbiota of LNM and NLNM groups formed two distinct clusters (see Fig.1B-C), indicating that there are certain compositional differences between the two groups. This finding motivated us to delve deeper into the key microbial species that cause these differences and assess their potential as biomarkers for LNM.

Through LEfSe analysis, we identified 21 gut microbiota associated with LNM. Notably, at the species level, we found that *Bacteroides plebeius* was enriched in the gut microbiota of LNM (see Fig.2B). *B. plebeius*, belonging to the *Bacteroidetes* phylum, is a bacterium successfully isolated from the human gut that consumes seaweed^[23]^. Compared with the gut microbiota of healthy adults^[22]^, the abundance of *B. plebeius* is significantly increased in the gut of CRC patients and those with adenomatous polyps. Further research investigating the composition of the gut microbiota during the transformation of colorectal adenomas to cancer revealed the importance of ten key species, including *Butyricimonas synergistica*, *Agrobacterium larrymoorei*, and *B. plebeius*^[24]^. In the analysis of the gut microbiota of Lynch syndrome patients prone to colorectal cancer^[25]^, *B. plebeius* showed overproliferation in the gut. These findings suggest that this bacterium may play a promotional role in the occurrence and development of colorectal cancer. In seaweed-treated mice, *B. plebeius* binds to IgA, which can limit bacterial growth through the chaining and aggregation of dividing cells^[26]^, a phenomenon consistent with our study (Fig.2C). Although it has been speculated that it may accelerate the progression of solid tumors by suppressing the body’s antitumor immune response^[27]^, to date, there have been no in-depth studies directly targeting the specific mechanisms between *B. plebeius* and the development of CRC. Overall, as a relatively novel bacterium in CRC research, *B. plebeius* is still in the early stages of study and requires further exploration. It is worth noting that in our predictive model construction process, *B. plebeius* ranked first in the gut microbiota feature ranking, and previous studies as well as our own research have shown that it is significantly enriched in the gut of CRC patients. This finding suggests that *B. plebeius* has the potential to become an important biomarker for identifying whether CRC patients have lymph node metastasis.

In order to identify the enriched biological pathways associated with the gut microbiome in relation to lymph node status in CRC, we utilized the PICRUSt2 software to predict the differential KEGG pathways between the gut microbiomes of NLNM and LNM-CRC patients. Our findings revealed that endocytosis is significantly enriched in NLNM group. Endocytosis is a core process by which cells acquire necessary substances and foreign entities from their external environment, and it encompasses a wide range, including nutrient particles, dietary components, bacterial cells, and toxic molecules^[28]^. In immune defense, macrophages can precisely use receptor-triggered endocytic pathways^[29]^ to internalize and degrade bacterial pathogens while presenting their antigens to stimulate the body’s immune response. Moreover, endocytosis also facilitates the entry of beneficial molecules released by commensal bacteria, such as bifidobacteria (e.g., polyamines) into intestinal epithelial cells^[30]^, exerting anti-inflammatory^[31]^ and antioxidant effects, strengthening the intestinal barrier, maintaining microbial balance, and playing a positive role in preventing CRC^[32]^. Researchers have also explored new strategies for delivering targeted drugs through endocytic pathways^[33]^, designing vitamin B12 conjugates using the overexpressed vitamin B12 receptor in colon cancer cells, and entering cancer cells precisely through receptor-mediated endocytosis, thereby inhibiting tumor growth and demonstrating the potential application value of endocytosis in disease treatment. Overall, the upregulation of endocytosis in NLNM-CRC patients may be the reason why their tumor progression is not obvious and difficult to metastasize distantly.

Moving forward, to clarify the relationship between the gut microbiota and the host immune system in CRC, we analyzed the correlation between LNM-associated differential gut microbiota and immune cells, as well as immune-related genes. *B. plebeius* caught our attention as it showed a positive correlation with resting NK cells (as shown in Fig.4C) and with the chemokine CXCL8 (Fig.5D). NK cells, as key cytotoxic lymphocytes, can precisely identify and eliminate target cells, release cytokines, enhance antiviral and immune responses^[34, 35]^. In the tumor environment, they directly attack cancer cells and coordinate with other immune networks to suppress tumor growth and spread^[36]^. Notably, NK cells have also shown potential therapeutic value in limiting the risk of liver metastasis^[37]^ in CRC patients. However, NK cells are not always activated; when they are in a resting state, their surveillance function is weakened, allowing tumor cells opportunities to escape and proliferate. Studies have shown that in high-risk colon tumor immune-infiltrated environments, resting NK cells and regulatory T cells (Tregs) accumulate significantly^[38]^. Our independent research also found that resting NK cells are enriched in LNM-CRC patients, and the aforementioned findings suggest that there may be some connection between resting NK cells and CRC lymph node metastasis.

Chemokines and cytokines are key regulators in tumor progression^[39]^, among which the CXCL12-CXCR4 signaling axis^[40]^ is particularly important for cancer metastasis. Upon activation, this axis enhances the transfer of microRNAs carried by CRC cell-derived extracellular vesicles to tumor-associated macrophages (TAM). These microRNAs modulate the function of the pre-metastatic niche (PMN) in TAM, promoting immune suppression, angiogenesis, and the formation of a pre-metastatic environment, thereby accelerating CRC growth and spread^[41]^. CXCL8, also known as IL-8, is an important chemokine whose role in CRC distant metastasis has been extensively studied. Angiogenesis is key to CRC progression, and CXCL8 promotes CRC angiogenesis through the CXCR2 receptor, providing survival resources^[42]^ for the tumor and creating conditions for metastasis. CXCL8 can also act in concert with CCL20 to synergistically induce epithelial-mesenchymal transition (EMT) through the PI3K/AKT-ERK1/2 signaling axis, thereby promoting CRC metastasis progression^[43]^. In addition, studies have found that *Streptococcus gallolyticus* can stimulate colon cancer cells to release IL-8^[44]^, while *Fusobacterium nucleatum* induces the secretion of IL-8 and CXCL1 by host cell binding and invasion^[45]^, promoting the migration of colorectal cancer cells and highlighting the synergistic role of the gut microbiota and chemokines in CRC. The upregulation of IL-8 in CRC epithelial cells has also been confirmed^[46]^. Combined with our correlation analysis, it can be speculated that the potential interaction between *B. plebeius* and IL-8 may be one of the possible reasons for CRC to undergo lymphatic metastasis and further distant metastasis.

In the analysis of the differential biological pathways between two groups of patients, we also found that *B. plebeius* is significantly positively correlated with the GOBP: Fructose metabolic process (r=0.84, p=0.004) (Fig.6C). As a monosaccharide, the metabolism and utilization of fructose in the gut are influenced by the gut microbiota, which in turn also affects the gut microbiota. Moderate intake of fructose can provide nutrition for beneficial gut bacteria, enhance the function of the intestinal barrier, regulate immune function, etc. However, excessive intake of fructose may alter the gut microbiota and disrupt the integrity of the intestinal epithelial barrier^[47]^, leading to health issues such as intestinal inflammation and cancer^[48]^. Bu et al., through in-depth analysis of CRC liver metastasis samples and in vivo metastasis models, revealed the important role of fructose metabolism in the metastatic process of CRC^[49]^. They found that with the progression of CRC, glycolysis, gluconeogenesis, and fructose metabolism all showed an upward trend. In particular, metastatic cancer cells in the liver significantly enhanced their fructose metabolic capacity by upregulating the expression of aldolase B, providing necessary energy support for the central carbon metabolism during tumor cell proliferation and further promoting the liver metastasis of CRC. In addition, Shen et al. also pointed out^[50]^ that under conditions of limited glucose supply, HCT116 and HT29 can increase the uptake and metabolism of fructose by upregulating the expression of the GLUT5 transporter protein, in order to maintain their core carbon metabolism activities and ensure the continuous growth and proliferation of tumor cells. These findings emphasize the potential role of fructose in the progression of CRC, especially in organs such as the liver and intestine that are directly involved in fructose metabolism. Cancer cells can flexibly adjust their metabolic mechanisms to utilize fructose to meet their metabolic needs. Therefore, according to our analysis, *B. plebeius*, which is significantly enriched in the gut of LNM-CRC patients, may provide the necessary energy for cancer cells to grow and metastasize by promoting the fructose metabolic process.

In the CRC clinical staging, LNM serves as a critical indicator in defining stage III disease, and its treatment plan significantly differs from that of the early stages of the disease. Given the importance of lymph node status in formulating treatment strategies, the scientific community is actively seeking more accurate and efficient predictive models. Notably, although many studies have focused on optimizing predictive models, research utilizing the gut microbiota, a key factor of the internal environment, to predict the LNM status in CRC is still scarce. Based on this finding, we used the significant characteristic differences in the gut microbiota of LNM-CRC and NLNM-CRC as fecal microbial markers to construct two advanced predictive models, RF and MLP, aiming to accurately distinguish between LNM-CRC and NLNM-CRC patients. The experimental results show that both the RF and MLP models achieved an area under the receiver operating characteristic curve well above 0.9 on the training set, and they also maintained a high AUC value close to 0.8 on the validation set, fully demonstrating the outstanding efficacy and stability of these two models in predicting the LNM status of CRC patients. This finding provides a low-cost, easy-to-implement, and highly efficient new strategy for predicting LNM in clinical practice and also opens up new horizons for personalized treatment of CRC.

The study also has its limitations: First, the collection of fecal samples is influenced by diet, digestion speed, and intestinal transit, and changes such as microbial DNA degradation, overgrowth, and death of certain species during storage may lead to data bias, making it difficult to fully reflect the changes in the gut microbiota. Second, the study only compared the fecal microbiota of CRC patients with negative and positive lymph nodes, lacking data from non-cancer patients and a comparison of mucosal and fecal microbiota. In the future, we plan to increase the collection of samples from both CRC lesions and adjacent normal mucosa to deepen the analysis of microbial communities, with the hope of accurately parsing the differences in the microbiome between diseased and normal tissues, thereby aiding in disease research.

## 4. Conclusion

In CRC patients, we identified 21 gut microbiota significantly associated with LNM, particularly *Bacteroides plebeius*, which was significantly enriched in LNM group and closely related to LNM-CRC. This species may play a key role in accelerating the progression of CRC and promoting lymphatic and distant metastasis. This process may involve an increase in resting NK cells, and the upregulation of expression of CXCL8 and fructose metabolic pathways. Furthermore, based on the characteristics of LNM-associated gut microbiota, the RF and MLP models we constructed showed significant clinical application prospects in predicting the LNM status of CRC patients.

## 5. Materials and Methods

### 5.1 Subject information and sample collection

With formal approval from the Medical Ethics Committee of the Guangxi Medical University Cancer Hospital, and based on the fully informed consent obtained from the patients, we successfully collected fecal samples from 236 CRC patients treated at the hospital from January 1, 2021, to December 31, 2021. After strict screening, 198 samples met the quality requirements for 16S rRNA sequencing. Subsequently, we meticulously extracted key demographic and clinical pathological characteristics from the medical records of these patients, including gender, age, TNM staging, tumor size, and microsatellite stability status. Specifically, in this study, we conducted an in-depth analysis of 147 fecal samples carrying information on lymph node metastasis status, specifically distinguishing 69 NLNM cases from 78 LNM cases. In addition, we carefully collected postoperative tumor samples from nine participants for further transcriptome sequencing analysis.

The strict criteria for sample inclusion ensured the quality and reliability of the research data, specifically including: (1). Patients who underwent surgery and received a clear pathological staging according to the 8th edition of the AJCC CRC staging criteria, or those diagnosed with colorectal adenocarcinoma through colonoscopic biopsy; (2). No history of other malignant tumors; (3). No history of intestinal diseases or acute complications such as intestinal perforation, intestinal obstruction, pelvic abscess, etc.; (4). No antitumor treatment of any kind received prior to fecal sample collection; (5). No use of antibiotics or gut microbiota modulators within one month prior to sample collection; (6). No history of consciousness disorders or other cognitive function impairments.

The fecal sample collection was carried out on the first day of the patient’s admission. We specifically instructed patients to collect the middle section of the stool using a sterile collection tube to prevent the mixture of urine. After collection, the samples were quickly aliquoted into 2 mL EP tubes (each containing approximately 200 milligrams) and immediately placed in a −80°C sterile ice box for long-term freezing preservation. At the same time, the freshly resected tumor tissue (about the size of a soybean, with a diameter of 3-5 millimeters) was properly stored in liquid nitrogen within 30 minutes of acquisition to maintain its biological activity and integrity.

### 5.2 16S rRNA sequencing and gut microbiota analysis

Before formal sequencing, we first conducted a thorough assessment of the DNA quality in the fecal samples. Using MOBIO PowerSoil DNA Extraction Kit, 200 milligrams of fecal samples were mixed with Tris-EDTA buffer to optimize the DNA extraction process. Subsequently, only high-quality DNA samples were selected for PCR amplification to ensure the accuracy of subsequent analyses. During the amplification process, we used specific primers 341F (5′-CCTACGGGNGGCWGCAG-3′) and 805R (5′-GACTACHVGGGTATCTAATCC-3′), which precisely target the V3 and V4 regions of the 16S rRNA gene, achieving selective amplification of the target fragments^[51]^. The amplified products were then identified by 2% agarose gel electrophoresis, focusing on bands in the range of 300 to 350 bp to ensure the accuracy and specificity of the amplified fragments. Using the Quant-iT PicoGreen dsDNA assay kit, we accurately determined the concentration of the PCR products, and all samples were mixed at equimolar concentrations. Subsequently, the combined samples were quantified a second time using the KAPA Library Quantification Kit KK4824 to ensure consistency of the samples before sequencing.

On the Illumina PE250 platform, we conducted high-throughput sequencing of qualified libraries using 2 × 250 bp chemistry reagents. After obtaining the raw sequencing data in FASTQ format, we utilized QIIME2 (Quantitative Insights Into Microbial Ecology version 2) to perform rigorous data filtering, eliminating low-quality sequences and retaining high-quality data for subsequent analysis.

Using the Greengene database v13.8, we annotated the gut microbiota, and subsequently performed ASV (Amplicon Sequence Variants)/OTU (Operational Taxonomic Units) extraction and analysis in the phyloseq package v1.26.1. We employed α-diversity indices (such as Chao1, ACE, Shannon, and Simpson indices) to assess the richness, diversity, and evenness of the gut microbiota. Venn analysis was utilized to characterize the similarities and differences in the composition of gut microbes among various samples, and this analysis was implemented through the vegan package v2.5.6.

To further explore the classification and comparative information within high-dimensional data, we employed the "mixOmics" package v6.6.2 to perform Partial Least Squares Discriminant Analysis (PLS-DA). Concurrently, we utilized the LEfSe software v1.0.0 to screen out gut microbial communities that significantly differentiate between lymph node metastasis and non-lymph node metastasis CRC patients under a strict threshold of LDA score |LDA score| > 2 and P value < 0.05. The influence of these differentially abundant microbial communities was assessed through LDA scores and, after logarithmic transformation, presented in an intuitive bar chart format using the ggplot2 package v3.4.0.

For the identified key gut microbial communities, we further utilized the PICRUSt2 software v2.3.0 to predict the potential enrichment of KEGG pathways between LNM-CRC and NLNM-CRC based on 16S rRNA sequencing data. By employing the non-parametric Mann-Whitney test, we compared the differences in α-diversity indices between the two groups and, with the aid of the vegan package v2.5.6, conducted an in-depth analysis of the enrichment of KEGG pathways between the groups. All data analyses were performed in the R software v4.3.2 environment, with statistical significance set at a P value < 0.05 (two-tailed test).

### 5.3 Transcriptome sequencing

RNA was extracted from tissue samples of nine CRC tumors using the Trizol® Total RNA Extraction Kit, four of which were from patients with lymph node metastasis and five from patients without lymph node metastasis. The integrity of the isolated RNA was verified by electrophoresis, and the purity was assessed using a mini UV-spectrophotometer. After removing rRNA, cDNA libraries were prepared according to the guidelines of the RNA-seq Sample Preparation Kit (VAHTS ™ Stranded mRNA-seq Library Prep Kit for Illumina®). These transcriptome libraries were sequenced on the Illumina NovaSeq 6000 system, generating 6G of data per sample. The quality of the sequencing data was evaluated using FastQC. Subsequently, the obtained sequences were aligned with the reference genome (hg38 version) using HISAT2 for gene expression analysis. StringTie was used to quantify the gene expression levels based on established gene models, and the expression abundance of each gene was calculated in terms of Transcripts Per Million (TPM).

### 5.4 Analysis of the CRC tumor immune microenvironment

CIBERSORT is an algorithm for quantifying the composition of immune cells from RNA sequencing data, relying on the unique gene expression signatures of different immune cells and employing a machine learning algorithm to identify and classify these gene expression profiles^[52]^. We used the CIBERSORT R version 1.03, in conjunction with known reference gene expression profiles and gene expression data from the composite samples to be analyzed, to create a model using support vector regression. The TPM matrix derived from the transcriptome sequencing was transformed into a matrix representing the relative proportions and functional states of 22 different immune cell types. The microbial composition matrix and the immune cell proportion matrix were integrated, and the correlation coefficients between each column of the merged matrix were calculated using the rcorr function in R.

### 5.5 Functional enrichment analysis of LNM-related transcriptome sequencing

In the R software v4.3.2 environment, we first conducted an in-depth analysis of RNA sequencing data by applying the ssGSEA (Single Sample Gene Set Enrichment Analysis) algorithm to gene sets in GMT format (downloaded from the GSEA website: https://www.gsea-msigdb.org/gsea/index.jsp, including c2.cp.kegg.v2022.1.Hs.symbols.gmt and c5.go.v2022.1.Hs.symbols.gmt). This algorithm calculates ssGSEA scores for each gene set in a sample based on the descending order of gene expression levels, quantifying the extent of coordinated upregulation or downregulation of members of a particular gene set in the sample. Subsequently, we utilized the "ssgsea" function in the "GSVA" package v1.46.0 to generate a gene set score matrix for each sample. To identify differences between lymph node metastasis and non-lymph node metastasis, we set the non-lymph node metastasis group as the control group and employed the "limma" algorithm integrated in the "TCGAbiolinks" package v2.25.3 to perform a differential analysis of GO items and KEGG pathways between the two groups, with the selection criteria set to P value < 0.05 and |log2FC| > 0. The GO analysis covered three aspects: biological process (BP), molecular function (MF), and cellular component (CC). To visually present these significantly different GO terms and KEGG pathways, we used the powerful plotting capabilities of the "ggplot2" package v3.4.0 to create volcano plots, effectively visualizing the analysis results.

### 5.6 Machine learning model building and identification of gut microbiota markers

We designed and implemented two machine learning architectures, Random Forest (RF) and Multilayer Perceptron (MLP), aimed at predicting LNM status of CRC patients based on gut microbiota features. RF is an ensemble machine learning method based on decision trees^[53]^, which improves the overall model’s predictive accuracy and stability by constructing multiple decision trees and combining their predictions. Its inherent feature selection mechanism is particularly powerful, and there have been studies exploring strategies to further optimize RF performance by reducing the feature space, and these efforts have successfully improved the model’s predictive accuracy in some scenarios^[54]^. RF not only occupies an important position in the modern field of machine learning but is also considered one of the cutting-edge technologies for building predictive models, showing a significant advantage in predictive performance compared to traditional methods^[55-57]^. MLP is a neural network-based machine learning model that performs high-level abstraction and classification of input data through multiple layers of nonlinear transformations. MLP typically consists of an input layer, one or more hidden layers, and an output layer, each containing multiple neurons that are interconnected through weights and biases^[58]^. MLP is not only short in training time and easy to implement but also provides a more robust model^[59]^, and it can solve more problems than a single-layer perceptron model. Therefore, MLP has shown unique advantages and potential in solving complex prediction tasks.

Using Python in conjunction with the SciKit Learn 0.18 library(https://scikit-learn.org/stable/), we set up a machine learning modeling environment and configured the model parameters by default. Based on 147 CRC patient gut microbiome data labeled with LNM, we randomly divided the dataset into training and testing parts in a 7:3 ratio. With the 21 types of gut bacteria as features, we selected species that are significantly related to LNM and ranked in the top 62% for importance, and built RF and MLP models based on this. We evaluated the model’s predictive performance on both the training and validation sets using ROC curves and AUC values, ensuring comprehensiveness and accuracy in our assessment.

### 5.7. Statistical methods

In R software version 4.3.2, we conducted statistical assessments using the Pearson chi-square test for categorical data and t-tests for continuous data related to clinical baseline characteristics. To establish correlations between the gut microbiome, immune cell counts, and immune-related gene expression, we performed Pearson correlation analysis using the Hmisc package version 5.1.1. The ggcorrplot software package version 0.1.4.1 was utilized for Spearman correlation analysis to examine the relationships between the dominant microbial groups and the enriched biological processes (BP), molecular functions (MF), and KEGG pathways. We employed various tools for visualizing the analysis results: pheatmap version 1.0.12 for heat maps, ggcorrplot package version 0.1.4.1 for correlation plots, ggplot2 version 3.4.0 for volcano plots, Igraph package version 1.6.0 for network graphs, and Cytoscape software version 3.10.1 for complex network analysis.

## Ethics approval and consent to participate

This study was approved by the Ethics and Human Subject Committee of Guangxi Medical University Cancer Hospital

## Author Contributions

Yongzhi Wu, Xiaoliang Huang, Zhen Wang, Zigui Huang, Jungang Liu, Weizhong Tang, Xianwei Mo: conceived and designed the experiments; Jungang Liu, Xiaoliang Huang, Yongzhi Wu, Yongqi Huang, Zigui Huang, Fuhai He, Binzhe Tang, Chuanbin Chen, Mingjian Qin, Xianwei Mo, Chenyan Long, Weizhong Tang: analyzed the data; Jungang Liu, Xiaoliang Huang, Yongzhi Wu, Yongqi Huang, Zigui Huang, Zhen Wang, Mingjian Qin, Fuhai He, Chuanbin Chen, Xianwei Mo, Chenyan Long, Binzhe Tang, Weizhong Tang: helped with reagents/materials/analysis tools; Jungang Liu, Xiaoliang Huang, Yongzhi Wu, Yongqi Huang, Chuanbin Chen, Zigui Huang, Mingjian Qin, Chenyan Long, Xianwei Mo, Binzhe Tang, Weizhong Tang: contributed to the writing of the manuscript. All authors reviewed the manuscript.

## Foundation

This work was supported by the Natural Science Foundation of Guangxi Province (Guangxi Natural Science Foundation) (2021GXNSFAA196008), Youth Science Foundation of Guangxi Medical University (GXMUYSF202402), Youth Research Project of Guangxi Medical University Affiliated Cancer Hospital (yuanqingji2023-10hao), Middle-aged and Young Teachers’ Basic Ability Promotion Project of Guangxi (Basic Ability Promotion Project of Guangxi) (2021KY0087), China Postdoctoral Science Foundation(2023MD734155) and Youth Science Foundation of Guangxi Medical University (GXMUYSF202357).

## Data Availability Statement

The original contributions presented in the study are included in the article material, further inquiries can be directed to the corresponding authors.

## References

[1] Bray F A-O, Laversanne M, Sung H A-O, et al.(2024) Global cancer statistics 2022: Globocan estimates of incidence and mortality worldwide for 36 cancers in 185 countries [J]. CA Cancer J Clin, 1542-4863, 74(73):229-263.doi:10.3322/caac.21834

[2] Sobhani I, Bergsten E, Couffin S, et al.(2019) Colorectal cancer-associated microbiota contributes to oncogenic epigenetic signatures [J]. Proc Natl Acad Sci U S A, 1091-6490 116(148):24285-24295.doi:10.1073/pnas.1912129116

[3] Han B, Zheng R, Zeng H, et al.(2024) Cancer incidence and mortality in China, 2022 [J]. J Natl Cancer Cent, 2667-0054 4(1):47–53.doi:10.1016/j.jncc.2024.01.006

[4] Imai H, Sawada K Fau - Sato A, Sato A Fau - Nishi K, et al.(2015) [Complete resection of liver metastases of colorectal cancer after high efficacy bevacizumab, S-1, and Cpt-11 combination chemotherapy] [J]. Gan To Kagaku Ryoho, 0385-0684, 42(41):101-104

[5] Rebersek M A-O.(2021) Gut microbiome and its role in colorectal cancer [J]. Bmc Cancer, 1471-2407, 21(21):1325.doi:10.1186/s12885-021-09054-2

[6] Holmes E, Li Jv Fau - Marchesi J R, Marchesi Jr Fau - Nicholson J K, et al.(2012) Gut microbiota composition and activity in relation to host metabolic phenotype and disease risk [J]. Cell Metab, 1932-7420, 16(15):559-564.doi:10.1016/j.cmet.2012.10.007

[7] Tilg H, Adolph T E, Gerner R R, et al.(2018) The Intestinal Microbiota in Colorectal Cancer [J]. Cancer Cell, 1878-3686, 33(36):954-964.doi:10.1016/j.ccell.2018.03.004

[8] Ranjbar M, Salehi R, Haghjooy Javanmard S, et al.(2021) The dysbiosis signature of Fusobacterium nucleatum in colorectal cancer-cause or consequences? A systematic review [J]. Cancer Cell Int, 1475-2867, 21(21):194.doi:10.1186/s12935-021-01886-z

[9] Wu Q L, Fang X T, Wan X X, et al.(2024) Fusobacterium nucleatum-induced imbalance in microbiome-derived butyric acid levels promotes the occurrence and development of colorectal cancer [J]. World J Gastroenterol, 2219-2840, 30(14):2018-2037.doi:10.3748/wjg.v30.i14.2018

[10] Markowiak-Kopeć P, Śliżewska K.(2020) The Effect of Probiotics on the Production of Short-Chain Fatty Acids by Human Intestinal Microbiome. Lid - 10.3390/nu12041107 [doi] Lid - 1107 [J]. Nutrients, 2072-6643, 12(14):1107.doi:10.3390/nu12041107

[11] Mcquade J L, Daniel C R, Helmink B A, et al.(2019) Modulating the microbiome to improve therapeutic response in cancer [J]. Lancet Oncol, 1474-5488, 20(22):e77-e91.doi:10.1016/S1470-2045(18)30952-5

[12] Jin M, Frankel W L.(2018) Lymph Node Metastasis in Colorectal Cancer [J]. Surg Oncol Clin N Am, 1558-5042, 27(22):401-412.doi:10.1016/j.soc.2017.11.011

[13] Förster R, Braun A Fau - Worbs T, Worbs T.(2012) Lymph node homing of T cells and dendritic cells via afferent lymphatics [J]. Trends Immunol, 1471-4981, 33(36):271-280.doi:10.1016/j.it.2012.02.007

[14] Mescoli C, Albertoni L Fau - Pucciarelli S, Pucciarelli S Fau - Giacomelli L, et al.(2012) Isolated tumor cells in regional lymph nodes as relapse predictors in stage I and Ii colorectal cancer [J]. J Clin Oncol, 1527-7755, 30(39):965-971.doi:10.1200/JCO.2011.35.9539

[15] Dai X, Dai Z, Fu J, et al.(2024) Prognostic significance of negative lymph node count in microsatellite instability-high colorectal cancer [J]. World J Surg Oncol, 1477-7819, 22(21):186.doi:10.1186/s12957-024-03469-4

[16] Compton C C, Greene F L.(2004) The staging of colorectal cancer: 2004 and beyond [J]. CA Cancer J Clin, 0007-9235, 54(56):295-308.doi:10.3322/canjclin.54.6.295

[17] Sleeman J P, Cady B Fau - Pantel K, Pantel K.(2012) The connectivity of lymphogenous and hematogenous tumor cell dissemination: biological insights and clinical implications [J]. Clin Exp Metastasis, 1573-7276, 29(27):737-746.doi:10.1007/s10585-012-9489-x

[18] Naxerova K A-O, Reiter J A-O, Brachtel E, et al.(2017) Origins of lymphatic and distant metastases in human colorectal cancer [J]. Science, 1095-9203, 357(6346):6355-6360.doi:10.1126/science.aai8515

[19] Kudo S E, Ichimasa K, Villard B, et al.(2021) Artificial Intelligence System to Determine Risk of T1 Colorectal Cancer Metastasis to Lymph Node [J]. Gastroenterology, 1528-0012, 160(164):1075-1084.e1072.doi:10.1053/j.gastro.2020.09.027

[20] Xia W A-O, Li D, He W A-O, et al.(2024) Multicenter Evaluation of a Weakly Supervised Deep Learning Model for Lymph Node Diagnosis in Rectal Cancer at Mri [J]. Radiol Artif Intell, 2638-6100, 6(2):e230152.doi:10.1148/ryai.230152

[21] Bremnes R M, AL-Shibli K Fau - Donnem T, Donnem T Fau - Sirera R, et al.(2011) The role of tumor-infiltrating immune cells and chronic inflammation at the tumor site on cancer development, progression, and prognosis: emphasis on non-small cell lung cancer [J]. J Thorac Oncol, 1556-1380, 6(4):824-833.doi:10.1097/JTO.0b013e3182037b76

[22] Senthakumaran T A-O, Moen A E F, Tannæs T M, et al. (2023) Microbial dynamics with Crc progression: a study of the mucosal microbiota at multiple sites in cancers, adenomatous polyps, and healthy controls [J]. Eur J Clin Microbiol Infect Dis, 1435-4373, 42(43):305-322.doi:10.1007/s10096-023-04551-7

[23] Kitahara M, Sakamoto M, Ike M, et al.(2005) Bacteroides plebeius sp. nov. and Bacteroides coprocola sp. nov., isolated from human faeces [J]. Int J Syst Evol Microbiol, 1466-5026, 55(Pt 55):2143-2147.doi:10.1099/ijs.0.63788-0

[24] Lin C A-O, Li B A-O X, Tu C A-O, et al.(2022) Correlations between Intestinal Microbiota and Clinical Characteristics in Colorectal Adenoma/Carcinoma [J]. Biomed Res Int, 2314-6141, 2022:3140070.doi:10.1155/2022/3140070

[25] Mori G, Orena B S, Cultrera I, et al.(2019) Gut Microbiota Analysis in Postoperative Lynch Syndrome Patients [J]. Front Microbiol, 1664-302X, 10:1746.doi:10.3389/fmicb.2019.01746

[26] Kearney S M, Gibbons S M, Erdman S E, et al.(2018) Orthogonal Dietary Niche Enables Reversible Engraftment of a Gut Bacterial Commensal [J]. Cell Rep, 2211-1247, 24(27):1842-1851.doi:10.1016/j.celrep.2018.07.032

[27] Kim Y, Kim G, Kim S, et al.(2024) Fecal microbiota transplantation improves anti-PD-1 inhibitor efficacy in unresectable or metastatic solid cancers refractory to anti-PD-1 inhibitor [J]. Cell Host Microbe, 1934-6069, 32(38):1380-1393.e1389.doi:10.1016/j.chom.2024.06.010

[28] Konishi H, Fujiya M Fau - Kohgo Y, Kohgo Y.(2015) Host-microbe interactions via membrane transport systems [J]. Environ Microbiol, 1462-2920, 17(14):931-937.doi:10.1111/1462-2920.12632

[29] Downey G P, Botelho Rj Fau - Butler J R, Butler Jr Fau - Moltyaner Y, et al.(1999) Phagosomal maturation, acidification, and inhibition of bacterial growth in nonphagocytic cells transfected with Fcgammariia receptors [J]. J Biol Chem, 0021-9258, 274(240):28436-28444.doi:10.1074/jbc.274.40.28436

[30] Uemura T, Stringer De Fau - Blohm-Mangone K A, Blohm-Mangone Ka Fau - Gerner E W, et al.(2010) Polyamine transport is mediated by both endocytic and solute carrier transport mechanisms in the gastrointestinal tract [J]. Am J Physiol Gastrointest Liver Physiol, 1522-1547, 299(292):G517-222.doi:10.1152/ajpgi.00169.2010

[31] Obayashi M, Matsui-Yuasa I Fau - Matsumoto T, Matsumoto T Fau - Kitano A, et al.(1992) Polyamine metabolism in colonic mucosa from patients with ulcerative colitis [J]. Am J Gastroenterol, 0002-9270, 87(86):736-740

[32] Goodwin A C, Destefano Shields Ce Fau - Wu S, Wu S Fau – Huso D L, et al.(2011) Polyamine catabolism contributes to enterotoxigenic Bacteroides fragilis-induced colon tumorigenesis [J]. Proc Natl Acad Sci U S A, 1091-6490, 108(137):15354-15359.doi:10.1073/pnas.1010203108

[33] Gupta Y, Kohli Dv Fau - Jain S K, Jain S K.(2008) Vitamin B12-mediated transport: a potential tool for tumor targeting of antineoplastic drugs and imaging agents [J]. Crit Rev Ther Drug Carrier Syst, 0743-4863, 25(24):347-379.doi:10.1615/critrevtherdrugcarriersyst.v25.i4.20

[34] Björkström N A-O X, Strunz B, Ljunggren H G.(2022) Natural killer cells in antiviral immunity [J]. Nat Rev Immunol, 1474-1741, 22(22):112-123.doi:10.1038/s41577-021-00558-3

[35] Cózar B, Greppi M A-O X, Carpentier S, et al. (2021) Tumor-Infiltrating Natural Killer Cells [J]. Cancer Discov, 2159-8290, 11(11):34-44.doi:10.1158/2159-8290.CD-20-0655

[36] Malmberg K J, Carlsten M, Björklund A, et al.(2017) Natural killer cell-mediated immunosurveillance of human cancer [J]. Semin Immunol, 1096-3618, 31:20-29.doi:10.1016/j.smim.2017.08.002

[37] Harmon C A-O X, Robinson M A-O, Hand F, et al. (2019) Lactate-Mediated Acidification of Tumor Microenvironment Induces Apoptosis of Liver-Resident Nk Cells in Colorectal Liver Metastasis [J]. Cancer Immunol Res, 2326-6074, 7(2):335-346.doi:10.1158/2326-6066.CIR-18-0481

[38] Zhang X, Zhao H, Shi X, et al.(2020) Identification and validation of an immune-related gene signature predictive of overall survival in colon cancer [J]. Aging (Albany NY), 1945-4589, 12(24):26095-26120.doi:10.18632/aging.202317

[39] Verbeke H, Struyf S Fau - Laureys G, Laureys G Fau – Van Damme J, et al.(2011) The expression and role of Cxc chemokines in colorectal cancer [J]. Cytokine Growth Factor Rev, 1879-0305, 22(25-26):345-358.doi:10.1016/j.cytogfr.2011.09.002

[40] Müller A, Homey B Fau - Soto H, Soto H Fau - Ge N, et al.(2001) Involvement of chemokine receptors in breast cancer metastasis [J]. Nature, 0028-0836, 410(6824):6850-6826.doi:10.1038/35065016

[41] Bhat A A, Nisar S, Singh M, et al.(2022) Cytokine- and chemokine-induced inflammatory colorectal tumor microenvironment: Emerging avenue for targeted therapy [J]. Cancer Commun (Lond), 2523-3548, 42(48):689-715.doi:10.1002/cac2.12295

[42] Watanabe K, Shiga K, Maeda A, et al.(2022) Chitinase 3-like 1 secreted from cancer-associated fibroblasts promotes tumor angiogenesis via interleukin-8 secretion in colorectal cancer. Lid - 3 [pii] Lid - 10.3892/ijo.2021.5293 [doi] [J]. Int J Oncol, 1791-2423, 60(61):63.doi:10.3892/ijo.2021.5293

[43] Cheng X S, Li Y F, Tan J, et al.(2014) CCL20 and CXCL8 synergize to promote progression and poor survival outcome in patients with colorectal cancer by collaborative induction of the epithelial-mesenchymal transition [J]. Cancer Lett, 1872-7980, 348(341-342):377-387.doi:10.1016/j.canlet.2014.03.008

[44] Abdulamir A S, Hafidh Rr Fau - Mahdi L K, Mahdi Lk Fau - AL-Jeboori T, et al.(2009) Investigation into the controversial association of Streptococcus gallolyticus with colorectal cancer and adenoma [J]. Bmc Cancer, 1471-2407, 9:403.doi:10.1186/1471-2407-9-403

[45] Casasanta M A-O, Yoo C A-O, Udayasuryan B A-O, et al.(2020) Fusobacterium nucleatum host-cell binding and invasion induces IL-8 and CXCL1 secretion that drives colorectal cancer cell migration. Lid - 10.1126/scisignal.aba9157 [doi] Lid - eaba9157 [J]. Sci Signal, 1937-9145, 13(641):eaba9157.doi:10.1126/scisignal.aba9157

[46] Cui G, Li G, Pang Z, et al.(2022) The presentation and regulation of the IL-8 network in the epithelial cancer stem-like cell niche in patients with colorectal cancer [J]. Biomed Pharmacother, 1950-6007, 152:113252.doi:10.1016/j.biopha.2022.113252

[47] Sellmann C, Priebs J, Landmann M, et al.(2015) Diets rich in fructose, fat or fructose and fat alter intestinal barrier function and lead to the development of nonalcoholic fatty liver disease over time [J]. J Nutr Biochem, 1873-4847, 26(11):1183-1192.doi:10.1016/j.jnutbio.2015.05.011

[48] Goncalves M A-O, Lu C A-O, Tutnauer J, et al.(2019) High-fructose corn syrup enhances intestinal tumor growth in mice [J]. Science, 1095-9203, 363(6433):1345-1349.doi:10.1126/science.aat8515

[49] Bu P, Chen K Y, Xiang K, et al.(2018) Aldolase B-Mediated Fructose Metabolism Drives Metabolic Reprogramming of Colon Cancer Liver Metastasis [J]. Cell Metab, 1932-7420, 27(26):1249-1262.e1244.doi:10.1016/j.cmet.2018.04.003

[50] Shen Z, Li Z, Liu Y, et al.(2022) GLUT5-Khk axis-mediated fructose metabolism drives proliferation and chemotherapy resistance of colorectal cancer [J]. Cancer Lett, 1872-7980, 534:215617.doi:10.1016/j.canlet.2022.215617

[51] Kozich J J, Westcott Sl Fau - Baxter N T, Baxter Nt Fau - Highlander S K, et al.(2013) Development of a dual-index sequencing strategy and curation pipeline for analyzing amplicon sequence data on the MiSeq Illumina sequencing platform [J]. Appl Environ Microbiol, 1098-5336, 79(17):5112-5120.doi:10.1128/AEM.01043-13

[52] Chen B, Khodadoust M S, Liu C L, et al.(2018) Profiling Tumor Infiltrating Immune Cells with Cibersort [J]. Methods Mol Biol, 1940-6029, 2018:1711:2243-2259.doi:10.1007/978-1-4939-7493-1_12

[53] Cutler A, Stevens J R.(2006) Random forests for microarrays [J]. Methods Enzymol, 0076-6879, 2006:2411:2422-2032.doi:10.1016/S0076-6879(06)11023-X

[54] Dimitriadis S I, Liparas D.(2018) How random is the random forest? Random forest algorithm on the service of structural imaging biomarkers for Alzheimer’s disease: from Alzheimer’s disease neuroimaging initiative (ADNI) database [J]. Neural Regen Res, 1673-5374, 13(16):962-970.doi:10.4103/1673-5374.233433

[55] Hu J, Szymczak S.(2023) A review on longitudinal data analysis with random forest. Lid - 10.1093/bib/bbad002 [doi] Lid - bbad002 [J]. Brief Bioinform, 1477-4054, 24(22):bbad002.doi:10.1093/bib/bbad002

[56] Mangino A A-O, Finch W A-O.(2021) Prediction With Mixed Effects Models: A Monte Carlo Simulation Study [J]. Educ Psychol Meas, 1552-3888, 81(86):1118-1142.doi:10.1177/0013164421992818

[57] Carracedo-Reboredo P, Liñares-Blanco J, Rodríguez-Fernández N, et al.(2021) A review on machine learning approaches and trends in drug discovery [J]. Comput Struct Biotechnol J, 2001-0370, 19:4538-4558.doi:10.1016/j.csbj.2021.08.011

[58] Haghighat F.(2021) Predicting the trend of indicators related to Covid-19 using the combined MLP-Mc model [J]. Chaos Solitons Fractals, 0960-0779, 152:111399.doi:10.1016/j.chaos.2021.111399

[59] Car Z, Baressi Šegota S A-O, Anđelić N, et al.(2020) Modeling the Spread of COVID-19 Infection Using a Multilayer Perceptron [J]. Comput Math Methods Med, 1748-6718, 2020:5714714.doi:10.1155/2020/5714714

